# An HIV-1 broadly neutralizing antibody from a clade C infected pediatric elite neutralizer potently neutralizes the contemporaneous and autologous evolving viruses

**DOI:** 10.1101/403469

**Authors:** Sanjeev Kumar, Harekrushna Panda, Muzamil Ashraf Makhdoomi, Nitesh Mishra, Haaris Ahsan Safdari, Heena Aggarwal, Elluri Seetharami Reddy, Rakesh Lodha, Sushil Kumar Kabra, Anmol Chandele, Somnath Dutta, Kalpana Luthra

**Author notes:** **Corresponding Author’s mailing address**: Dept.of Biochemistry, All India Institute of Medical Sciences, Ansari Nagar, New Delhi-110029, and India. Equal contribution (Harekrushna Panda, Muzamil Ashraf Makhdoomi, Nitesh Mishra).

## Abstract

Broadly neutralizing antibodies (bNAbs) have demonstrated protective effects against HIV-1 in primate studies and recent human clinical trials. Elite-neutralizers are potential candidates for isolation of HIV-1 bNAbs and coexistence of bNAbs such as BG18 with neutralization susceptible autologous viruses in an HIV-1 infected adult elite controller has been suggested to control viremia. Disease progression is faster in HIV-1 infected children than adults. Plasma bNAbs with multiple epitope specificities are developed in HIV-1 chronically infected children with more potency and breadth than in adults. Therefore, we evaluated the specificity of plasma neutralizing antibodies of an antiretroviral naïve HIV-1 clade C chronically infected pediatric elite neutralizer AIIMS_330. The plasma antibodies showed broad and potent HIV-1 neutralizing activity with >87% (29/33) breadth, median inhibitory dilution (ID50) value of 1246 and presence of N160 and N332-supersite dependent HIV-1 bNAbs. The sorting of BG505.SOSIP.664.C2 T332N gp140 HIV-1 antigen-specific single B cells of AIIMS_330 resulted in the isolation of an HIV-1 N332-supersite dependent bNAb AIIMS-P01. The AIIMS-P01 neutralized 67% of HIV-1 cross-clade viruses; exhibited substantial indels despite limited somatic hypermutations; interacted with native-like HIV-1 trimer as observed in negative stain electron microscopy and demonstrated high binding affinity. In addition, AIIMS-P01 potently neutralized the coexisting and evolving autologous viruses suggesting the coexistence of vulnerable autologous viruses and HIV-1 bNAbs in AIIMS_330 pediatric elite neutralizer. Further studies on such pediatric elite-neutralizers and isolation of novel HIV-1 pediatric bNAbs may provide newer insights to guide vaccine design.

**Importance:** More than 50% of the HIV-1 infections globally are caused by clade C viruses. Till date, there is no effective vaccine to prevent HIV-1 infection. Based on the structural information of the currently available HIV-1 bNAbs, attempts are underway to design immunogens that can elicit correlates of protection upon vaccination. Here we report the isolation and characterization of an HIV-1 N332-supersite dependent bNAb AIIMS-P01 from a clade C chronically infected pediatric elite neutralizer. The N332-supersite is an important epitope and is one of the current HIV-1 vaccine targets. AIIMS-P01 potently neutralized the contemporaneous and autologous evolving viruses and exhibits substantial indels despite low somatic hypermutations. Taken together with the information on infant bNAbs, further isolation of bNAbs contributing to the plasma breadth in HIV-1 infected children may help to better understand their development and characteristics, which in turn may guide vaccine design.

## Introduction

The promising results of animal challenge studies and human clinical trials of potent HIV-1 broadly neutralizing antibodies (bNAbs) in inhibiting viral infection and recent FDA approval of Ibalizumab in combination with antiretroviral therapy for individuals with multidrug resistant HIV-1 has raised hopes for the development of effective HIV-1 therapeutic vaccines (1–4). Based on the structural information of the currently available HIV-1 bNAbs, attempts are underway to design immunogens that can elicit correlates of protection in the vaccinees %(5, 6). Studies conducted in chronically HIV-1 infected adult donors suggest that it takes at least 2 to 3 years of infection for the antibodies to undergo affinity maturation in response to viral escape, in order to develop potent HIV-1 bNAbs (7– 9). The HIV-1 bNAbs evolve in rare infected individuals who are antiretroviral naïve long-term non-progressors (LTNPs) or viremia controllers and the top 1% among them are classified as ‘elite neutralizers’ based on their plasma neutralizing activity (10). The HIV-1 bNAbs isolated from chronically infected adult elite neutralizers exhibit signature characteristic features like high somatic hypermutations (SHM), insertions or deletions (indels), long CDRH3 length, high potency and broad viral neutralization breadth (11–16). However, most adult bNAbs, with the exception of BG18 normally fail to neutralize coexisting autologous viruses (7, 12, 17–20), but no such data is available on pediatric elite neutralizer(s).

HIV-1 infection in children is mostly caused by vertical transmission (21). Early treatment with antiretroviral therapy (ART) delays disease progression, however, it fails to prevent HIV-1 infection (22). Without ART, disease progression is faster in infected children than in adults with median time to AIDS being one year, compared to ten years in ART naïve infected adults (23), due to the immaturity and limited diversity of the immune system (24– 26). Recently, plasma bNAbs have been shown to evolve in HIV-1 infected infants %(27, 28). Longitudinal studies revealed that children infected at birth, until 3 months post infection (pi) had high titers of NAbs effective against a few viruses, after which their titers decreased; passively transferred maternal NAbs probably being the source of the NAbs at this stage. The titers of potent NAb again increased with time, suggesting the development of de novo immune responses in all the infants about one year pi (28). This plasma mapping study showed that HIV-1 bNAbs can develop early in life and that functional B cells persist in these infants to produce bNAbs, irrespective of high viremia and faster disease progression than adults (28, 29). The high viral load, both in infants and adults possibly promotes the development of NAb breadth (28, 30–32). Moreover, BF520.1, one of the HIV-1 N332-supersite dependent bNAb isolated from an infant, one-year post-infection, has shown broad neutralizing activity, despite limited SHMs (33), with the absence of indels, unlike the bNAbs isolated from adults (13, 16) suggesting that infant bNAbs might be evolved by different pathways than adult bNAbs.

Few longitudinal studies, including ours, conducted on pediatric HIV-1 directed humoral immune response showed that plasma bNAbs develop in select chronically infected children and slow progressors, with diverse epitope specificities, (24, 34–37), and with higher breadth and potency than infected adults (35). Further, the bNAbs present in the plasma of infected children demonstrated similar epitope specificities as that observed for the adult bNAbs (24). Recently, from pooled peripheral blood mononuclear cells (PBMCs) of select HIV-1 clade C infected pediatric long-term non-progressors (LTNPs), we constructed an anti-HIV-1 human single chain recombinant fragment (scFv) phage library, from which one of the CD4-binding site (CD4bs) targeted scFv 2B10 demonstrated broad neutralizing activity with a breadth of 78% (38).

Herein, we have characterized the plasma neutralizing activity and epitope specificities of a chronically HIV-1C infected pediatric elite neutralizer AIIMS_330. HIV-1 specific single B cells from this donor were sorted using BG505.SOSIP.664.C2 T332N gp140 envelope trimer as antigenic bait followed by isolation of an N332-supersite dependent pediatric bNAb AIIMS-P01 that neutralized the autologous coexisting and evolving viruses.

## Results

### Identification of a rare pediatric elite neutralizer by longitudinal neutralization activity analysis of the plasma antibodies

The AIIMS_330 donor is an Indian HIV-1C chronically infected 9-year-old boy who had acquired the infection at birth through vertical transmission. At the time of sampling (2015), he was antiretroviral (ART) naïve, the CD4 T cell counts and plasma viral loads were 1033 cells/µl and 85,400 RNA copies/ml respectively. The plasma antibody neutralization activity of AIIMS_330 donor against a few HIV-1 viruses has been previously documented (37, 39, 40). In order to assess the evolution of antibody response in this donor, we further characterized AIIMS_330 (2015) plasma against a large panel of heterologous HIV-1 tier 1, 2 and 3 pseudoviruses (n=33) (Figure 1A, Table S1). The AIIMS_330 plasma antibodies showed broad and potent HIV-1 neutralizing activity with >87% (29/33) breadth, median inhibitory dilution (ID50) value of 1246 (Figure 1A, Table S1) and a neutralization score of 2.9 (10), thereby qualifying AIIMS_330 to be an “elite neutralizer”. The AIIMS_330 plasma showed V1V2 and V3-glycan directed neutralization specificities, identified using Env-pseudoviruses with point mutations at N160 (CAP45 clade C and BG505 clade A) and N332 (BG505 clade A and CAP256 clade C) glycans and their corresponding wild-type viruses (Figure 1B and 1C). However, no reduction in ELISA binding of plasma antibodies with CD4bs probe RSC3 mutant protein Δ371I/P363N, as compared to its wild-type protein RSC3 core, was observed. In addition, the plasma antibodies did not bind to MPER peptides either (Figure 1D), suggesting the absence of CD4bs and MPER directed antibodies in the 2015 time point AIIMS_330 plasma, as was also seen in its earlier 2013 time point plasma sample (38). Based on the elite neutralizing activity exhibited by AIIMS_330 (2015) plasma, this donor was selected for isolating anti-HIV-1 bNAbs.

**Figure 1:**
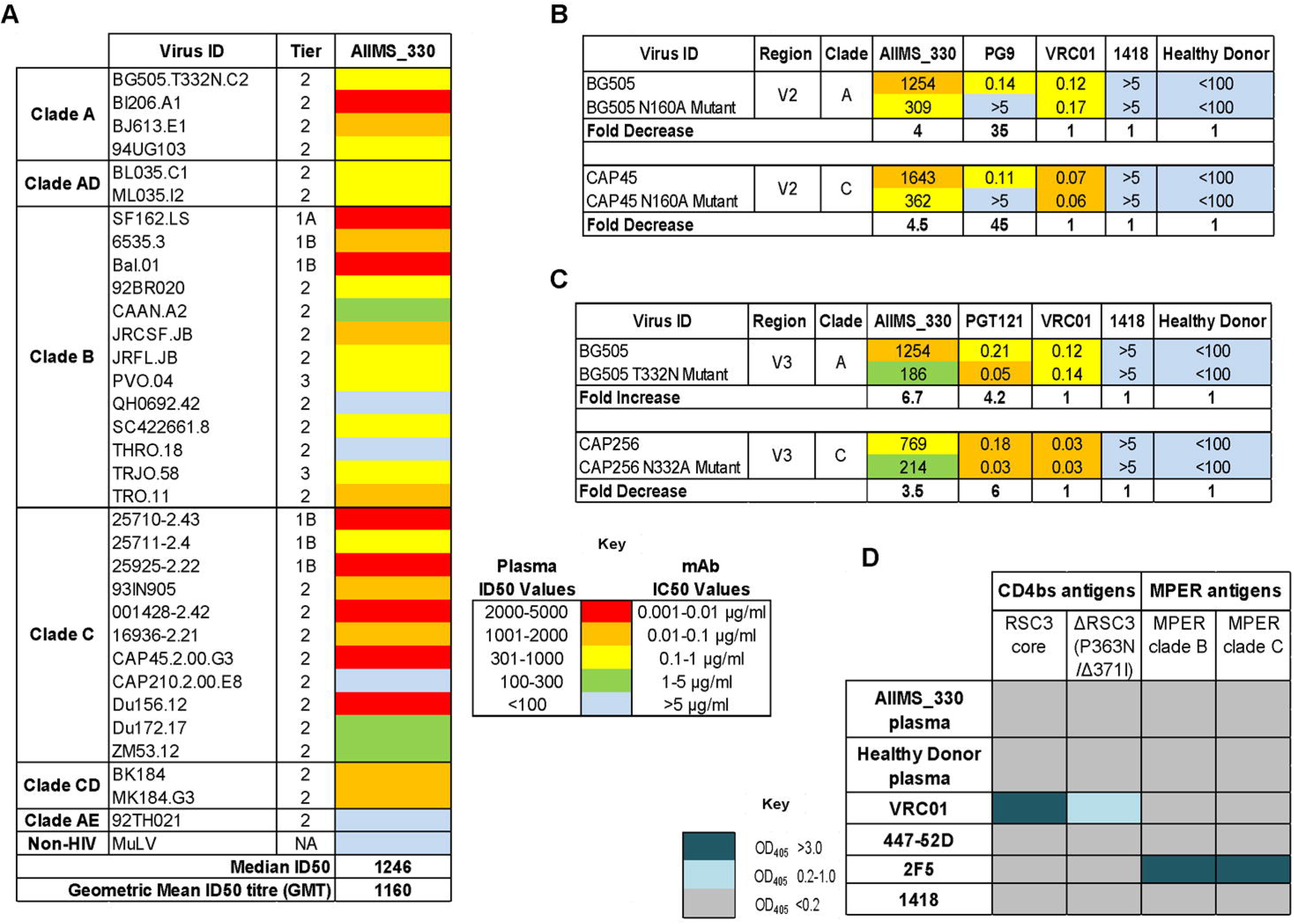
Plasma characterization identified AIIMS_330 as a pediatric elite neutralizer. Heat map showing the neutralization ID50 titers of AIIMS_330 plasma antibodies against a heterologous panel of HIV-1 viruses (n=33). MuLV was used as a negative virus control. Heat map showing the N160 dependent analysis of AIIMS_330 plasma using heterologous clade A virus BG505 and autologous clade C virus CAP45. Values are neutralization ID50 plasma titers or monoclonal antibodies IC50 titers. PG9 was a positive control for the V2 region, while VRC01 (CD4bs), 1418 (parvovirus mAb) and healthy donor plasma were used as negative controls. Reduction in ID50/IC50 titers of >3-fold with mutant as compared to wild-type virus was considered as positive dependence. (C) Heat map showing the N332 dependent analysis of AIIMS_330 plasma using heterologous clade A virus BG505 and autologous clade C virus CAP256. Values are neutralization ID50 plasma titers or monoclonal antibodies IC50 titers. PGT121 was a positive control for the V3-glycan N332 region. Reduction in ID50/IC50 titers of >3-fold with mutant as compared to wild-type virus was considered as positive dependence. (C) Heat map showing the ELISA results in OD at 405 nm to determine the CD4bs directed binding of AIIMS_330 plasma antibodies using RSC3 core protein, mutant protein RSC3 Δ371I/P363N and MPER binding using clade B and C 25 mer MPER peptides. VRC01 and 2F5 were used as positive controls for CD4bs and MPER respectively, whereas 447-52D (V3 region) and 1418 were used as a negative control.

### Pediatric antibody AIIMS-P01 demonstrated broad and potent HIV-1 neutralizing activity

To sort HIV-1 antigen-specific single B cells, initially the antigenic bait, avi-tagged trimeric BG505.SOSIP.664.C2 T332N gp140 protein was expressed, purified and biotinylated, as described previously (14), followed by its binding analysis with existing anti-HIV-1 bNAbs to confirm the binding reactivity of the biotinylated trimeric protein with the HIV-1 bNAbs and absence of binding with non-NAbs (Figure S1A and S1B). Then, a total of 22 BG505.SOSIP^+^ single B cells with the phenotype of CD3-, CD8-, CD14-, Aqua-(Dead cells), CD19+, IgG+, and IgM-were sorted from 2 million stored PBMCs of AIIMS_330 pediatric elite neutralizer (Figure 2A). Next, DNA was synthesized from single B cells. By nested PCR based amplification of antibody variable heavy and light chain genes using specific primers (41), we obtained seven pairs of heavy and light chain genes amplified from single B cell and sequenced. By applying the criteria of having SHM >5% (42) in the amplified antibody genes, we identified a singular pediatric HIV-1 bNAb AIIMS-P01. Therefore, for further experiments, only AIIMS-P01 was characterized. Next, to determine the neutralization potential of AIIMS-P01, its neutralization activity was tested at a concentration ranging from 50µg/ml to 0.001µg/ml, against a heterologous panel of HIV-1 viruses, using TZM-bl based neutralization assay (43) (Figure 2B, 2C and Table S2). The AIIMS-P01 bNAb effectively neutralized a panel of viruses comprising of susceptible tier-1 to highly resistant tier-3 viruses of different clades (Figure 2B and Table S2). Of the viruses tested, AIIMS-P01 neutralized 68% (20/29) clade C viruses and 78% (15/19) clade B viruses. Most of the viruses neutralized by AIIMS-P01 bNAb were also neutralized by the contemporaneous plasma antibodies, except for a tier-2 clade B virus QH0692 (Figure 2C). Three of the viruses TRJO, CAP45, and ZM53 were not neutralized by the AIIMS-P01 bNAb at the tested concentrations, although they were susceptible to neutralization by the contemporaneous plasma antibodies (Figure 2C). These results demonstrate that the broad viral neutralizing activity of AIIMS-P01 was comparable with that of the contemporaneous plasma antibodies. The pediatric bNAb AIIMS-P01 did not neutralize clade AE viruses (that do not have an Asn glycan at the 322 amino acid residue of the V3 loop), as has also been observed with the existing N332 dependent adults and infant HIV-1 bNAbs, suggesting their N332 glycan dependency (12, 13, 16, 33, 44). The overall breadth of AIIMS-P01 was >67% (neutralized 51/76 HIV-1 viruses), with a geometric mean IC50 titer of 0.5µg/ml (Figure 2B).

**Figure 2:**
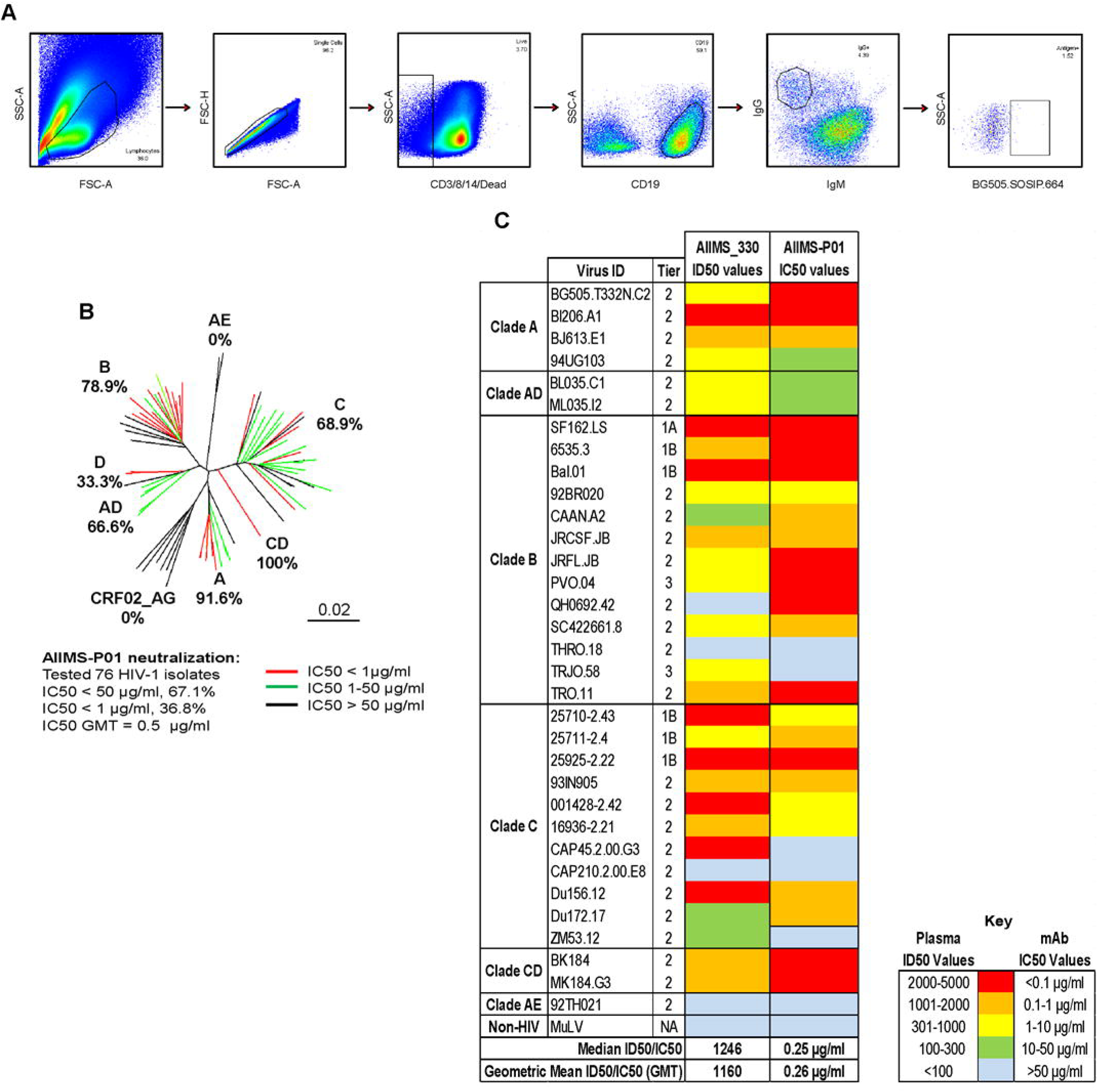
AIIMS-P01 demonstrated broad HIV-1 neutralizing activity and potently neutralized the contemporaneous and evolving viruses. (A) Single B cell sorting by FACS of AIIMS_330 pediatric elite neutralizer using BG505.SOSIP.664.C2 T332N gp140 envelope trimer as antigenic bait. (B) Dendrogram of neutralization IC50 values of AIIMS-P01 bNAb against a panel of heterologous HIV-1 viruses (n=76). (C) Heat map depicting AIIMS_330 plasma ID50 titers and IC50 values of contemporaneous AIIMS-P01 bNAb tested against viruses of different clades (n=33).

### AIIMS-P01 neutralized autologous contemporaneous and evolving viruses

To evaluate the ability of AIIMS-P01 to neutralize the autologous contemporaneous viruses of 2015 time point, HIV-1 rev/env cassette was amplified by single genome amplification (SGA) method as described previously (45), using cDNA as template synthesized from plasma viral RNA. The neutralization susceptibility of these autologous HIV-1 enveloped pseudoviruses to AIIMS-P01 was tested, along with the highly potent second-generation HIV-1 adult bNAbs, first generation NAbs and non-neutralizing antibodies (Figure 3A). The AIIMS-P01 potently neutralized all three autologous contemporaneous viruses designated as 330.15 series (Figure 3A). Two out of three contemporaneous viruses were also neutralized by adult bNAbs PGT121 (N332-directed) and 10E8 (MPER directed). Further, we generated the evolving 2016 autologous viruses (designated with 330.16 series) and tested their neutralization susceptibility to AIIMS-P01. We found that all of the evolving autologous viruses were neutralized by AIIMS-P01 (Figure 3A) and also by PGT121, 10E8, and PGT145 (V1V2 directed) adult bNAbs. These results suggest the coexistence of vulnerable autologous viruses and bNAbs in AIIMS_330 pediatric elite neutralizer, as has also been observed with BG18, isolated from an adult HIV-1 infected donor EB354 (12). Six out of the seven AIIMS_330 autologous viruses, were resistant to many of the potent adult bNAbs, except 330.16.E6 (Figure 3A). The phylogenetic analysis (Figure 3B) depicts the evolution of AIIMS_330 autologous viruses at different time points.

**Figure 3:**
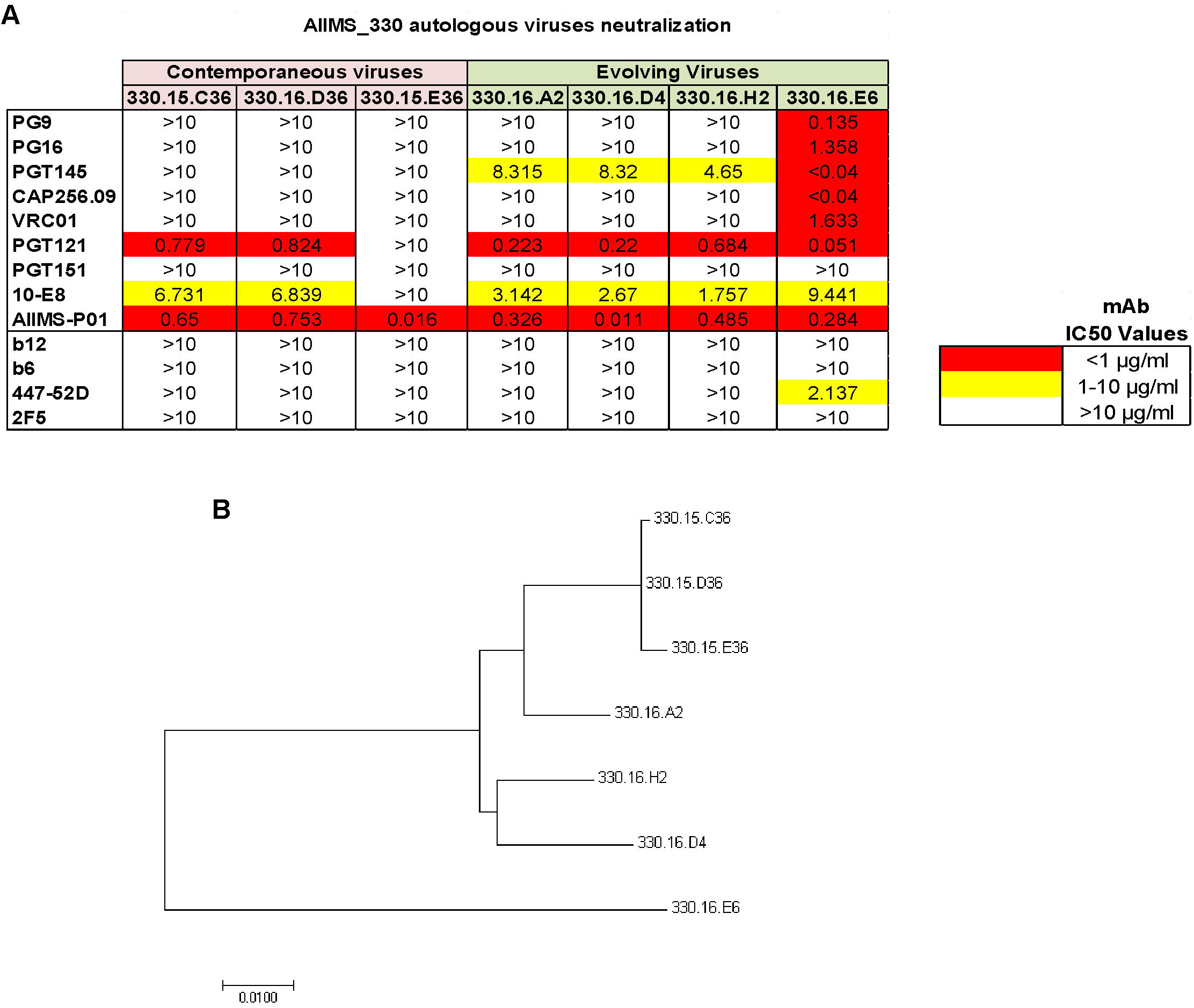
AIIMS-P01 neutralized autologous contemporaneous and evolving viruses. (A) Heat map of neutralization IC50 values of pediatric AIIMS-P01 bNAb and adults’ HIV-1 second-generation bNAbs (PG9, PG16, PGT145, CAP256.09, VRC01, PGT121, PGT151, 10E8), first generation NAbs (b12, 2F5 and 447-52D) and non-NAbs (b6) tested at 10µg/ml, against the autologous contemporaneous virus of AIIMS_330 (2015) depicted as (330.15) and successive time point (2016), depicted as (330.16). (B) Maximum-Likelihood phylogenetic tree analysis of AIIMS_330 autologous viruses of two time points constructed by MEGA7 software.

### AIIMS-P01 targeted the N332 supersite at the base of the V3 region of HIV-1 envelope

The absence of N332 glycan in clade AE viruses (44), furthered by the inability of AIIMS-P01 to neutralize clade AE viruses, suggests the N332 directed neutralizing dependency of this bNAb. In addition, a 55-fold reduction in IC50 neutralization activity of AIIMS-P01 against the clade C mutant CAP255.16 N332A, as compared to its wild-type CAP255.16 WT virus, confirmed its N332 directed epitope specificity. Furthermore, a 72-fold increase in IC50 was observed when this bNAb was tested against the BG505.C2 T332N mutant that contained the N332 glycan, as compared to its wild-type BG505.C2 WT that lacked a glycan in this position (Figure 4A). The N332 dependent neutralizing activity of AIIMS-P01 was further confirmed by competition ELISA using N332 dependent bNAbs 10-1074, PGT121 and BG18 (Figure 4B) (12, 13, 16). The binding of AIIMS-P01 with N332 region on the native-like BG505.SOSIP.664 trimeric envelope was observed in cell-surface binding assays (Figure 4C). These findings were further substantiated by negative staining electron microscopy (EM) (Figure 4D and 4E). The BG505.SOSIP.664.C2 T332N D-7324 tagged trimeric envelope glycoprotein was expressed, purified and its purity was assessed using blue native-PAGE (6) (Figure S2). Negative staining EM and reference free classification were performed to visualize the interaction of the native-like HIV-1 trimeric protein with AIIMS P01-Fab and AIIMS-P01 IgG respectively (Figure S3A, S3B, and S3C). Analysis of the complex of AIIMS-P01 Fab with BG505.SOSIP.664.C2 T332N gp140 HIV-1 trimeric antigen by negative stain EM confirmed that AIIMS-P01 showed binding reactivity with amino acid residues at the base of the V3 region (Figure 4D and 4E). The three Fab fragments were clearly visible in reference-free 2D class averages (Figure 4D and S5) and 3D EM envelope (Figure 4E and S4) suggesting the presence of three suitable epitopes in the HIV-1 trimeric protein that has to interact with Fab fragments. On the contrary, the density of AIIMS-P01 IgG with HIV-1 trimer in AIIMS P01-IgG complex is not clearly visible. The plausible explanation for this is that IgG density was averaged out during data processing owing to its inherent flexibility though it binds with HIV trimer. The three-dimensional (3D) reconstruction of HIV-1 trimer with AIIMS-P01 Fab at 26Å resolution (Figure 4E, S4A, S4B, and S4C) indicates that the AIIMS-P01 Fab interacts with the V3-region of the soluble HIV-1 trimer. Recent studies show that our structure is quite similar to other published structures of HIV trimers with PGV04 and PGT122 (46). Overall, the substitution of the Asn332 glycan with Ala resulted in a total reduction in neutralization activity of AIIMS-P01, whereas no reduction was observed with N160A, N156A mutants (Table S3), showing the absence of V1V2 directed neutralization activity of this bNAb. Moreover, no reduction in binding was observed between CD4bs probes RSC3 core and RSC3 mutant Δ37I/P363N (Table S3) demonstrating that pediatric bNAb AIIMS-P01 was dependent exclusively on HIV-1 V3-glycan N332.

**Figure 4:**
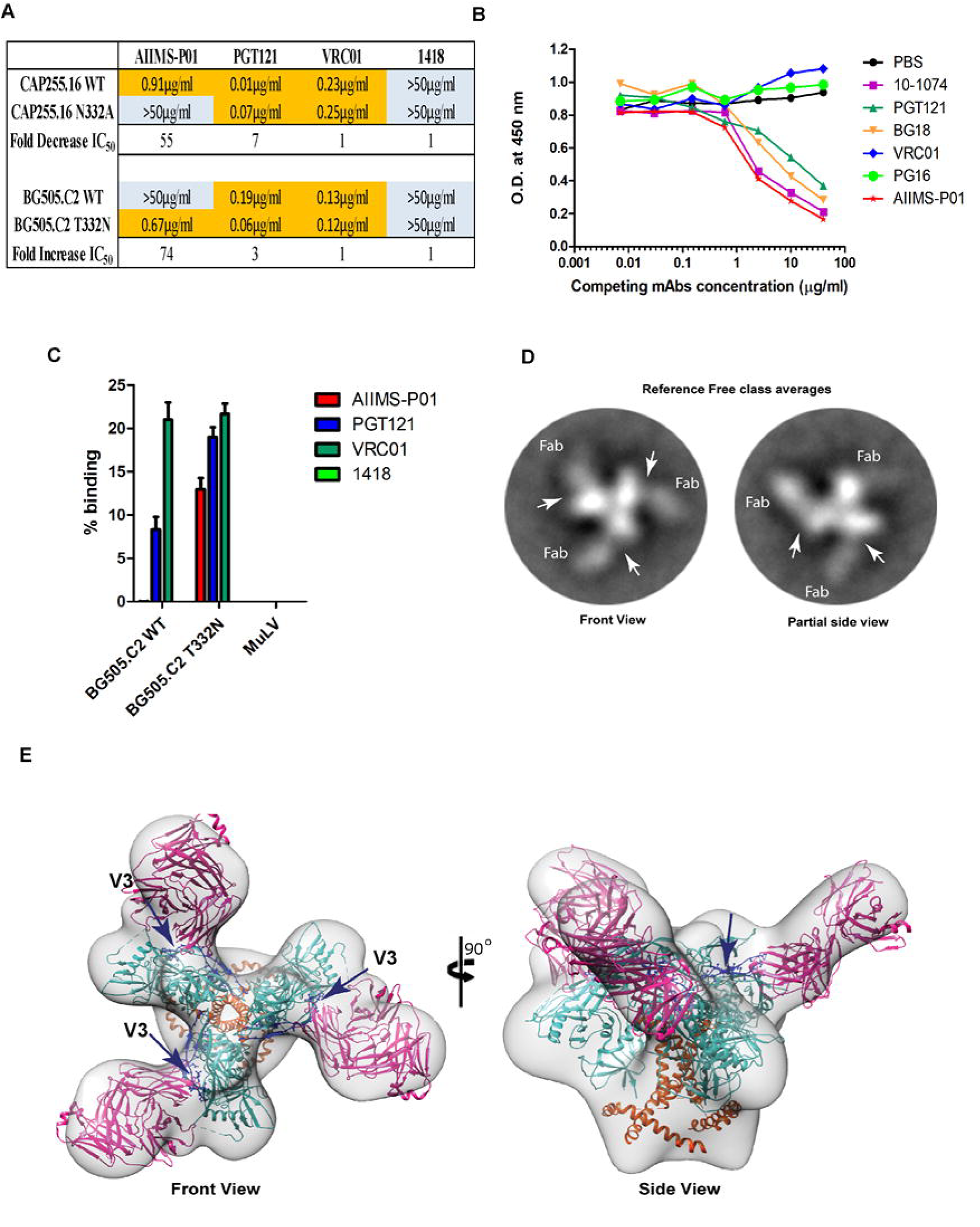
AIIMS-P01 targets the N332 supersite. (A) Neutralization IC50 values of monoclonal antibodies for their N332 dependence. PGT121 was used as positive control, VRC01 and 1418 were used as negative controls. (B) Competition ELISA findings: biotinylated AIIMS-P01 was added at a fixed concentration of 5µg/ml with increasing concentration of competing monoclonal antibodies. The black line indicates no competition in the absence of competing antibody. (C) Cell-surface binding assay results of the percentage binding of the monoclonal antibodies. Monoclonal antibodies VRC01, 1418 and MuLV virus were used as negative controls. PGT121 was used as positive control. (D) Reference-free 2D class averages of AIIMS-P01 Fabs bound with HIV-1 BG505.SOSIP.664 T332N gp140 trimer. The left average is the front view and the right average is the partial side view, which shows the density of AIIMS-P01 Fabs bound with HIV-1 BG505.SOSIP.664 T332N gp140 trimer. The arrows depict the V3 region of the HIV-1 trimer, which interacts with AIIMS-P01 Fab. (E) 3D-model reconstructed from the negative stain EM micrographs of AIIMS-P01 Fab in complex with BG505.SOSIP.664 T332N gp140 trimer. The crystal structure of HIV trimer (4NCO) and Fab (3PIQ) was docked into the EM envelope.

### AIIMS-P01 showed high affinity with native-like HIV-1 trimeric protein

Next, we determined the binding potential of AIIMS-P01 with heterologous HIV-1 monomeric clade C and B envelopes and observed comparable binding efficiency (Figure 5B, 5C, and 5D). In addition, the binding of AIIMS-P01 with native-like HIV-1 trimeric glycoprotein was comparable with that of PGT121 and PGT151 adults’ bNAbs (Figure 5A). ELISA binding showed that AIIMS-P01 displayed high binding with both trimeric HIV-1 antigen and heterologous monomeric antigens (Figure 5A, 5B, 5C, and 5D). To further validate these results, the affinity of the AIIMS-P01 with trimeric gp140 and monomeric gp120 antigens was assessed using Surface Plasmon Resonance (SPR). AIIMS-P01 displayed low off-rate and higher affinity (KD=9.8nM) with BG505.SOSIP.664 T332N in SPR experiments than monomeric BG505 gp120 protein (KD=50nM) (Figure 5E and 5F). These results demonstrate that AIIMS-P01 exhibit high affinity with the native-like HIV-1 trimer.

**Figure 5:**
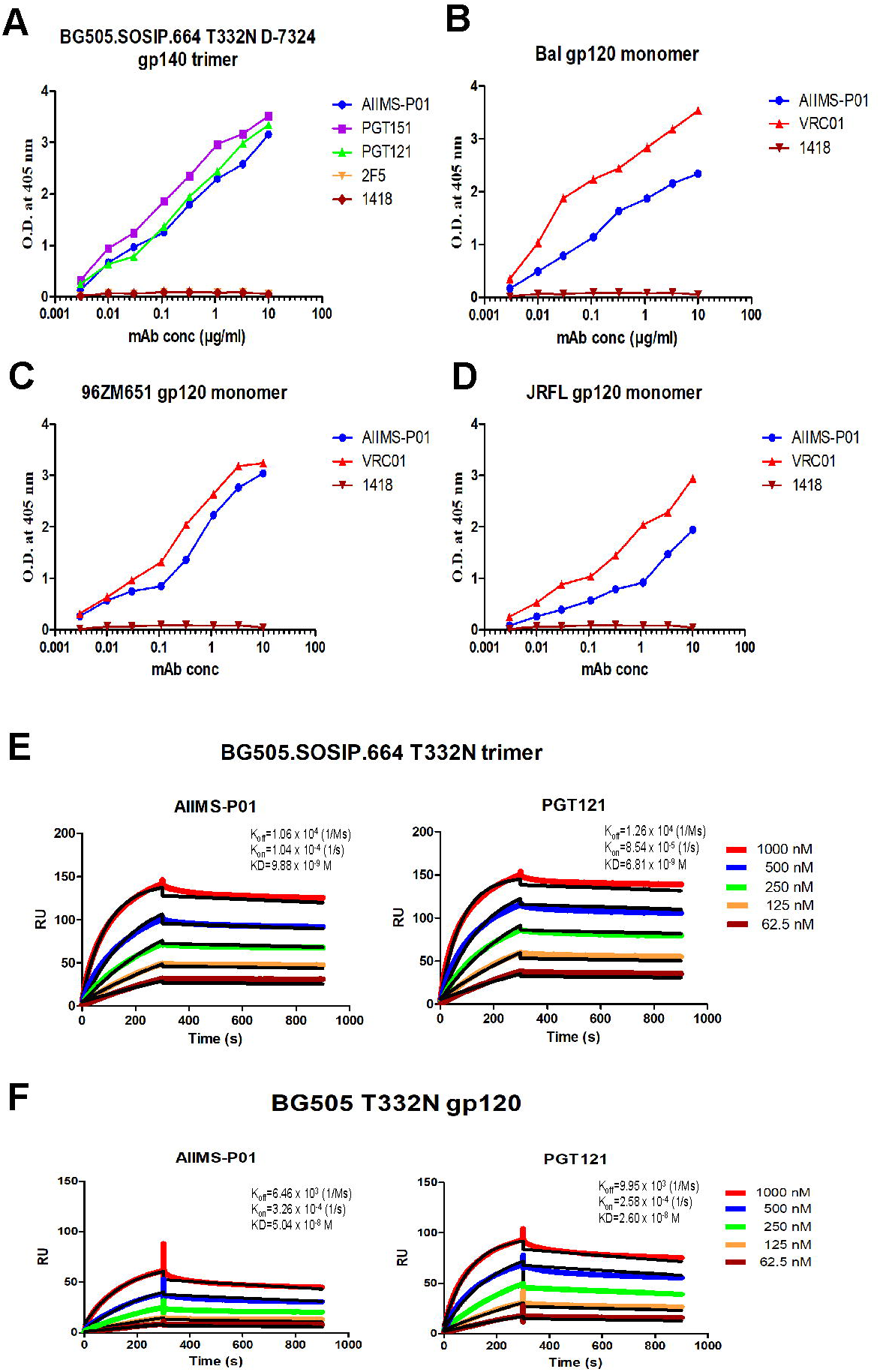
AIIMS-P01 showed strong binding with a native-like HIV-1 antigen. (A) Binding curves of AIIMS-P01 with BG505.SOSIP.664 T332N D-7324-tagged gp140 trimer. Here, PGT151 trimer specific HIV-1 bNAb and PG121 (N332) bNAb were used as positive controls, 2F5 (MPER), 1418 were used as negative controls. (B) (C) and (D) are binding curves of AIIMS-P01 with monomeric gp120 antigens. Here, VRC01 was used as positive controls 1418 was used as negative control. (E) SPR sensorgrams of AIIMS-P01 and PGT121 with BG505.SOSIP gp140 trimer. (F) SPR sensorgrams of AIIMS-P01 and PGT121 with BG505 gp120 monomer. Here, PGT121 was used positive control.

### AIIMS-P01 exhibited the presence of indels with limited somatic hypermutations

The sequence comparison of pediatric bNAb AIIMS-P01 with its respective germline heavy and light chain gene, showed the presence of indels (+5 amino acids insertions in heavy chain framework region 3 (FRH3)), that has not been observed in the heavy chain of the other N332 directed bNAbs (PGT121 and 10-1074) (Figure 6A) (13, 16). In addition, a limited frequency of SHM, 7% in the heavy chain and 5% in the light chain were found in AIIMS-P01 (Figure 6B). Further, AIIMS-P01 had similar germline gene family of IGVH4-59*01 as that of adult bNAbs of 10-1074 and the PGT121 lineage (13, 16), with a distinct light chain gene family of IGLV1-47*02 (Figure 6A and 6B). Similar to the N332 targeted adult bNAbs (12, 13, 16, 44) and infant bNAbs of BF520.1 class (33), the CDRH3 region of AIIMS-P01 was 19 amino acids long. Long CDRH3 has been shown to be important for interaction with HIV glycans (47, 48) (Figure 6B). Phylogenetic analysis of the N332 dependent AIIMS-P01 displayed a close relation with the N332 dependent adult bNAb 10-1074 (Figure 6C). In addition, a comparison of the neutralization activity of AIIMS-P01, adult and infant bNAbs of N332 epitope specificity revealed that AIIMS-P01 exhibited neutralization breadth comparable with that of the adult N332 bNAbs (Figure 6D). A comparison of the gene usage, CDR length, the presence of indels, epitope specificities and neutralizing activity between N332 directed bNAbs AIIMS-P01 and adult bNAbs, along with bNAbs of different specificities are shown in a table (Figure 6E).

**Figure 6:**
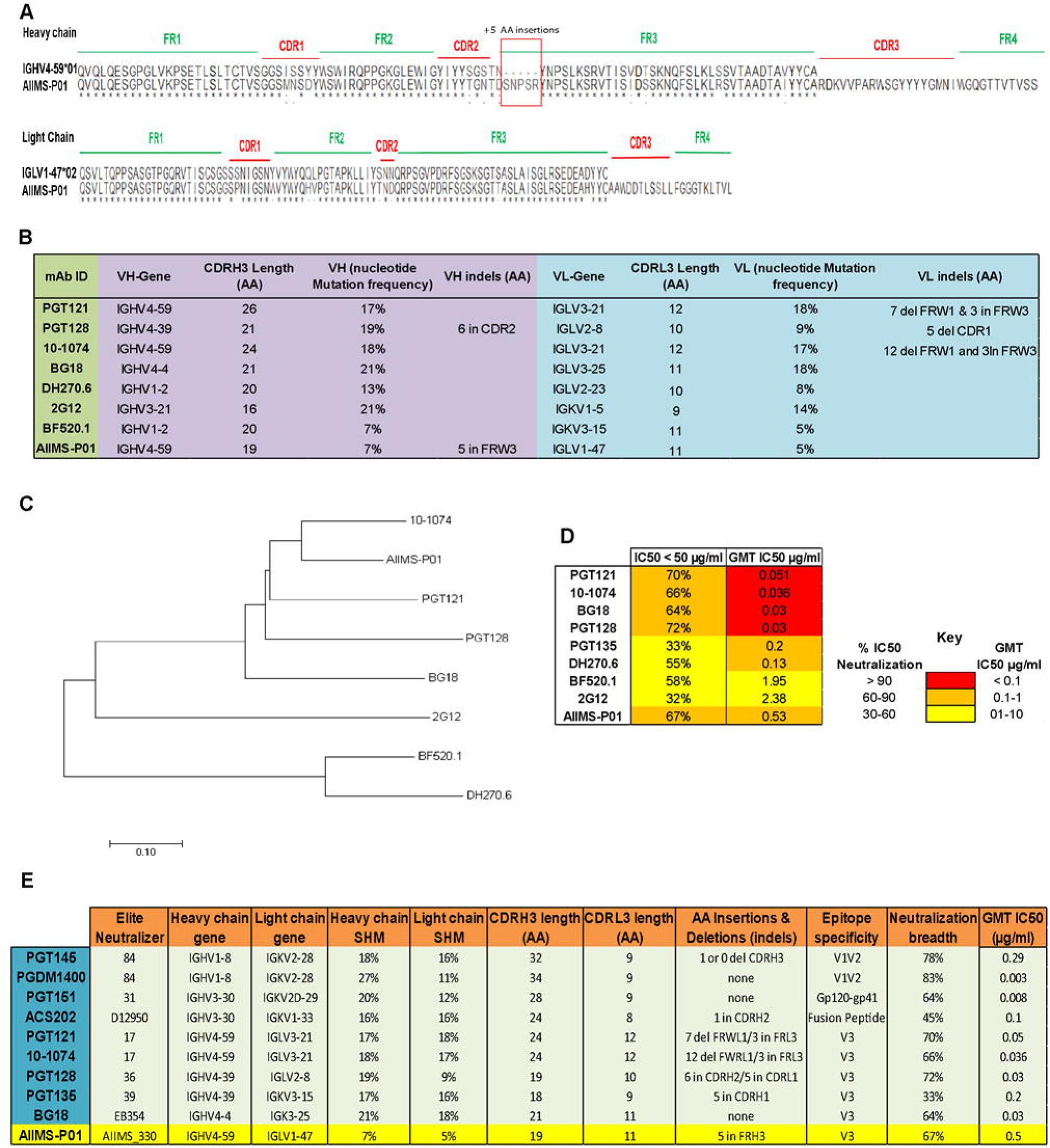
AIIMS-P01 exhibited the presence of indels with modest somatic hypermutations. (A) Alignment of heavy and light chain amino acid sequence with respective germline antibody gene sequences. Alignment was done using clustalX software and BioEdit software. (B) Image showing the AIIMS-P01 comparison with the characteristic features of N332 directed bNAbs of adults and an infant. Here, in, insertion and del, deletion. (C) Maximum likelihood tree for phylogenetic analysis of N332 directed bNAbs, which was made by Mega7 software. (D) Heat map showing the comparison of AIIMS-P01 neutralization IC50 values with adult and infant N332 dependent bNAbs. (E) This is a comparison of the characteristics of HIV-1 bNAbs of adult and pediatric origin identified from all the elite neutralizers reported so far. Here, in, insertion and del, deletion.

### AIIMS-P01 showed ADCC activity with absence of polyreactivity

The ADCC activity of the AIIMS-P01 and its contemporaneous plasma antibodies was tested by infecting target cells with NL4-3 and JRFL viruses. The MuLV virus was used as a negative control. Both AIIMS-P01 bNAb and its contemporaneous plasma antibodies showed ADCC activity (Figure 7A and 7B). The AIIMS-P01 bNAb did not show any polyreactivity when tested for binding with multiple antigens, suggesting the lack of binding to non-HIV-1 antigens and cellular self-antigens (Figure S6).

**Figure 7:**
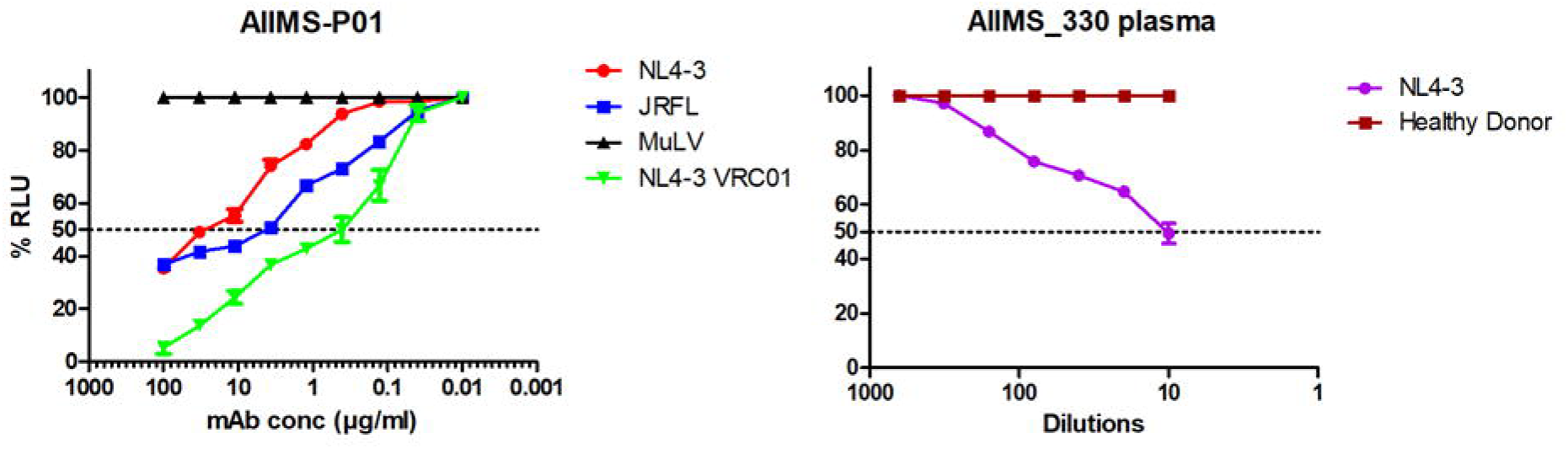
AIIMS-P01 exhibited ADCC activity. (A) ADCC activity analysis of AIIMS-P01 with NL4-3 and JRFL viruses. VRC01 was used as positive control and MuLV was used as a negative virus control. (B) The ADCC activity of AIIMS_330 plasma. The healthy donor was used as negative control.

## Discussion

Implementation of novel high-throughput techniques such as memory B cell culture and single B cell sorting using antigenic bait led to the successful isolation of second-generation potent HIV-1 human bNAbs (49). The epitopes defined by HIV-1 bNAbs form the basis for vaccine design using the Reverse Vaccinology 2.0 strategy (50). BNAbs such as BG18 can successfully neutralize the coexisting autologous viruses suggesting that similar bNAbs can be present in viremic controllers (12) Goo et al have for the first time shown the development of bNAbs in infants within one year of age (28). Further, the polyclonal antibodies with multiple specificities found in chronically infected children (24, 36) showed greater potency and breadth than the adult bNAbs (24, 35), providing an impetus for isolating bNAbs from such chronically infected children. The recent isolation of the infant bNAb BF520.1 (33), suggested the presence of precursor B cells in infants that can be stimulated by HIV-1. Mapping the specificities of bNAbs at different stages of infection; those elicited early in infection (28, 33), and ones found in chronically infected children (24, 35, 36), may provide useful information for combinatorial immunization strategies to elicit similar bNAbs in the vaccinees. Recently, we (36) and others (24, 35) observed the longitudinal evolution of plasma cross-neutralization antibody response with multiple epitope specificities in select antiretroviral naïve HIV-1C chronically infected children (24). It is pertinent to isolate bNAbs from infected individuals whose plasma antibodies have been extensively mapped for the presence of bNAbs, desirably the LTNPs and elite neutralizers (10, 50).

This study describes for the first time, a pediatric bNAb AIIMS-P01 IgG1, isolated from single B cell of an ART naïve HIV-1C chronically infected pediatric LTNP AIIMS_330, and identified as an elite neutralizer in this study, based on plasma mapping analysis. This AIIMS_330 donor belonged to a rare cohort of vertically transmitted ART naïve HIV-1C infected children that have been followed up for more than a decade (37–40). The AIIMS-P01 bNAb isolated from this elite neutralizer showed comparable neutralization efficiency against clade C and B HIV-1 viruses thereby suggesting that this bNAb may neutralize major viral clades responsible for infection globally.

The AIIMS-P01 bNAb is dependent exclusively on the N332 supersite epitope, similar to the HIV-1 bNAbs PGT121, 10-1074, DH270 and BG18 isolated from adult donors (12, 13, 16, 44) and BF520.1, an infant derived bNAb (33). The N332 supersite is an important epitope and is one of the current HIV-1 vaccine targets (5, 6). The frequency of somatic hypermutations (7%) observed in AIIMS-P01 was comparable to that found in the infant bNAb BF520.1 (33) and the bNAbs from adults (3.8% to 32.6%) (51, 52). Furthermore, AIIMS-P01 showed 5 amino acid (AA) insertions in the third framework region of the antibody heavy chain (FRH3), as has been observed to be prevalent in adult bNAbs, with in-frame indels ranging from 0 to 7 AA) (13, 15, 16, 53). The high frequency of indels in adult bNAbs has been shown to increase with and is predicted by the frequency of SHM (54), contributing to their high neutralization potency and breadth (47, 48, 51, 55, 56). However, it is difficult to elicit similar bNAbs with high SHM and indels by immunization (57), although such antibodies are generated in macaques, with an average SHM of 10% and with the presence of indels, on immunization with trimeric HIV-1 envelopes (58). It may be possible that AIIMS-P01 has attained high breadth by virtue of the substantial indels as reported earlier for N332 and CD4bs directed adults bNAbs that indels are critical for making contacts with protein or glycans on HIV (47, 48, 55, 56). The AIIMS-P01 bNAb has acquired indels with limited SHMs, whether such bNAbs can be elicited in vaccinees, needs to be addressed. AIIMS-P01 belongs to an IGVH4-59*01 germline gene family, similar to the existing bNAbs of PGT121 series and 10-1074, isolated from an adult elite neutralizer (13, 16). The sequence comparison, alignment and phylogenetic analysis of AIIMS-P01 with existing N332 glycan-dependent bNAbs, displayed a close relation to the 10-1074 class of adult bNAbs (13).

The neutralizing activity of AIIMS_330 plasma antibodies mapped to V1V2 and V3 glycan epitopes and corroborates with observations of multiple epitope specificities of the plasma antibodies found in chronically infected children (24, 36), supporting that bNAbs from pediatric elite neutralizers can be directed to similar neutralizing determinants as that found in adult elite neutralizers. Likewise, the IGVH1-2 antibody gene usage in infant bNAb BF520.1 (33) is also seen in DH270.6 bNAb from adults (44). Further, the binding interactions of the AIIMS-P01 Fab with the native-like BG505.SOSIP trimeric envelope glycoprotein, as observed in reference-free class averages and negative staining 3D reconstruction was possibly due to the long CDRH3 region of AIIMS-P01, that may promote contact with the envelope glycan residues (47, 48). AIIMS-P01 showed the ability to effectively neutralize not only the autologous contemporaneous viruses; but also autologous viral escape mutants of the successive time points, suggesting the coexistence of N332 dependent bNAbs and vulnerable viruses in AIIMS_330 pediatric elite neutralizer, as has been documented in EB354 elite controller, with the coexistence of BG18 bNAbs and susceptible contemporaneous autologous viruses (12). Apart from BG18 bNAb, isolated from an adult elite neutralizer, that neutralized autologous viruses (12), the current bNAbs do not neutralize contemporaneous autologous viruses (12, 17–20). The 3.5-6.7-fold ID50 increase in contemporaneous plasma neutralizing activity of N332 mutant pseudoviruses, in comparison to the 38-78-fold increase in AIIMS-P01 IC50 values with N332 mutant pseudoviruses of different clades, indicates that bNAbs of specificities other than N332 glycan are present in the plasma of this donor, may be N160 dependent, as observed in plasma mapping results, but, not captured by our antigen specific sorting method. Furthermore, AIIMS-P01 neutralization was comparable to the contemporaneous plasma neutralization breadth, suggesting N332 dependent AIIMS-P01 like bNAbs were substantially present in the contemporaneous AIIMS_330 plasma. The pediatric bNAb AIIMS-P01 showed high affinity with native-like soluble trimeric envelope and demonstrated heterologous HIV-1 clade neutralization activity. The inability to neutralize the viruses of AE clade, which lack the N332 glycan, further confirms its N332 neutralization dependence, as has also been seen earlier with other N332 dependent bNAbs (12, 13, 44). In addition, AIIMS-P01 showed effector functions and the absence of polyreactivity.

To the best of our knowledge, the present study for the first time reported a singular pediatric elite neutralizer followed by isolation and characterization of an HIV-1 N332-supersite dependent pediatric bNAb AIIMS-P01. The AIIMS-P01 contains indels with limited SHM and belongs to the antibody gene family of the PGT121 class of HIV-1 bNAbs. The AIIMS-P01 bNAb potently neutralized its coexisting and evolving autologous viruses. The immunotherapeutic potential of AIIMS-P01 may be tested alone and in combination with antiretroviral drugs for its ability to prevent and treat the HIV-1 infection. As this study was based on the single HIV-1 chronically infected child, our findings encourage to further isolate and characterize the HIV-1 bNAbs contributing to the plasma breadth in chronically infected children to better understand their development and characteristics which in turn may provide newer insights to guide immunogen design.

## Material and Methods

### Ethics statement and study subject

5ml of blood from HIV-1 clade C infected AIIMS_330 pediatric donor was drawn after the approval of this study by the institutional ethics committee, All India Institute of Medical Sciences (AIIMS), New Delhi, India (IEC/59/08.01.16 and IEC/NP-536/04.11.2013) and obtaining written informed signed consent form from the parents/guardian. The peripheral blood mononuclear cells (PBMCs) were isolated by ficoll density gradient method after separating the plasma by centrifuging at 360×g for 10 min.

### HIV-1 pseudoviruses generation

The HIV-1 pseudoviruses were produced in HEK 293T cells by co-transfecting the full HIV-1 gp160 envelope plasmid and a pSG3ΔEnv backbone plasmid. Briefly, 1×10^5^ cells in 2ml complete DMEM (10% fetal bovine serum (FBS) and 1% penicillin and streptomycin antibiotics) were seeded per well of a 6 well cell culture plate (Costar) the day prior to co-transfection for HIV-1 pseudovirus generation. For transfection, envelope (1.25µg) to delta envelope plasmid (2.50µg) ratio was 1:2, this complex was made in Opti-MEM (Gibco) with a final volume of 200µl for each well of the 6 well plate and incubated for 5 minutes at room temperature. Next, 3µl of PEI-Max transfection reagent (Polysciences) (1mg/ml) was added to this mixture, mixed well and further incubated for 15 min at room temperature. This mixture was then added dropwise to HEK 293T cells supplemented with fresh complete DMEM growth media and incubated at 37°C for 48 hours. Pseudoviruses were then harvested by filtering cell supernatants with 0.45 mm sterile filter (mdi) and stored frozen at −80°C as aliquots.

### HIV-1 neutralization assays

The HIV-1 neutralization assays of AIIMS-330 plasma antibodies and HIV-1 monoclonal antibodies (mAbs) were done as described earlier (43). Neutralization was measured as a reduction in luciferase gene expression after a single round of infection of TZM-bl cells (NIH AIDS Reagent Program) with HIV-1 envelope pseudoviruses. The TCID_50_ of the HIV-1 pseudovirus was calculated and 200 TCID_50_ of the virus was used in neutralization assays by incubating with 1:3 serially diluted heat-inactivated plasma at starting dilution of 1/100 for 1 hour at 37°C. After that, freshly trypsinized TZM-bl cells in growth medium (complete DMEM with 10% FBS and 1% penicillin and streptomycin antibiotics) containing 50μg/ml DEAE Dextran and 1 mM Indinavir (in case of primary isolates) at 10^5^ cells/well were added and plates were incubated at 37°C for 48 hours. Virus controls (cells with HIV-1 virus only) and cell controls (cells without virus and antibody or plasma) were included. MuLV was used as a negative control. After the incubation of the plates for 48 hours, luciferase activity was measured using the Bright-Glow Luciferase Assay System (Promega). ID50 values of the plasma sample and IC50 for antibodies were calculated. Values were derived from a dose-response curve fit with a non-linear function using the GraphPad Prism 5 software (San Diego, CA).

### Binding analysis of AIIMS_330 plasma antibodies

All proteins (RSC3 core and mutant) and peptides (MPER-B and C) were coated on ELISA plates at a concentration of 2μg/ml in 0.1M NaHCO_3_ (pH 9.6) and incubated overnight at 4°C. Next day, plates were blocked with 300μl/well of 15% FBS and 2% BSA in RPMI for 1.5 h, followed by addition of 100 μl/well of heat inactivated plasma at different dilutions (1:100, 1:300, 1:1,000, 1:3,000, and 1:10,000) or HIV-1 monoclonal antibodies (VRC01, 447-52D and 2F5) and anti-Parvovirus mAb 1418 each at 5-0.001 μg/ml concentrations and incubated for 1 hour at 37°C. Each of the above steps was followed by washing with 1×PBS (Phosphate Buffered Saline) with 0.1% Tween 20. Then, 100μl of alkaline phosphatase (AP)-conjugated anti-human IgG Fc antibody [1:2,000 diluted in 1×PBS (Southern Biotech)] was added to the plates and the immune complexes were reacted with AP substrate in 10% DAE buffer (1 mg/ml) and absorbance was read at 405 nm.

### Expression and purification of BG505.SOSIP.664 T332N trimeric proteins

The BG505.SOSIP.664.C2 T332N gp140 trimeric proteins with D-7324-tag and Avi-tag were expressed in HEK 293F cells and purified by methods described previously (6, 14). Purity was assessed by blue native polyacrylamide gel electrophoresis (BN-PAGE) and binding reactivity with HIV-1 bNAbs was assessed by ELISA. The presence of native-like SOSIP trimers with D-7325-tagged was further confirmed by negative-stain electron microscopy. The purified avi-tagged BG505.SOSIP.664.C2 T332N gp140 trimeric protein was first biotinylated using the BirA500 Ligase Reaction Kit (Avidity) according to the manufacturer’s protocol, biotinylation of trimer was assessed by ELISA as described earlier (14) and used for the sorting of single B cells of donor AIIMS_330.

### HIV-1 BG505.SOSIP trimer-specific single B cell sorting by FACS

The frozen PBMCs (2×10^7^ cells) of AIIMS_330 pediatric elite neutralizer were thawed in complete DMEM containing 50U/ml of benzonase enzyme (Novagen). Next, cells were stained using a panel of fluorophore-labeled antibodies directed to CD3 (V500), CD8 (V500), CD14 (V500), CD19 (PE-Cy7) human IgM (FITC) and human IgG (BV421) and Live/Dead Fixable aqua-fluorescent reactive dye (Invitrogen) (to exclude dead cells). All fluorochrome-cojugated antibodies were purchased from BD Biosciences. In addition, to sort antigen specific single B cells, the HIV-1 BG505.SOSIP.664 gp140 trimer was labelled with streptavidin-APC (Life Technologies) followed by its addition to the PBMCs at 4°C for 30 min, followed by twice washing with ice-cold 1×PBS. Cells with a phenotype of CD3-/CD8-/CD14-/CD19+/IgG+/IgM-/BG505.SOSIP.664 trimer APC+ were singly sorted on a BD FACS AriaIII sorter (BD Biosciences) into 96-well PCR plates (BioRad) containing 20 µl/well lysis buffer consisting of 1U/ml RNase OUT (Thermo Fisher), 0.3125% Igepal CA-630, 1×SuperScript III First-Strand Buffer and 6.25 mM dithiothreitol (DTT) provided with the Superscript III Reverse Transcriptase kit (Thermo Fisher). The sorting data was collected using FACSDiva software (BD Biosciences) and analyzed using FlowJo software (FlowJo, LLC).

### Single B cell RT-PCR, amplification of antibody genes and cloning

The amplification of variable heavy and light chain antibody genes was performed as described earlier (41, 53, 59) with few modifications. The frozen 96-well plate containing single B cell sorted was first thawed on ice followed by cDNA synthesis using 1µl of 200U/well Superscript III reverse transcriptase (Invitrogen), 1 µL of 10mM dNTP mix (Thermo Fisher), 0.5µL of 50ng/ml random hexamers (Thermo Fisher) and 0.5µL of OligodT (Thermo Fisher) under the following cycling parameters: 25°C 10 min, 42°C 10 min, 50°C 50 min, 55°C 10 min, 85°C for 5 min and hold at 4°C. Next, the first strand cDNA was amplified by two-step nested-PCR. In the first step, 0.4µL of the mixture of 5µM IgH, Ig? or Igλ chain specific primers were used for the amplification of antibody variable genes of heavy and light chains in a final reaction volume of 25µL using 0.3µL of DreamTaq DNA Polymerase (5U/µL) (Thermo Fisher), 3µL of first-strand cDNA, 0.3µL of 10mM dNTP mix (Thermo Fisher) under the following cycling parameters: 95°C 5 min; 35 cycles of 95°C 30 sec, 50 or 52°C 60 sec, 72°C 1 min, 72°C 10 min and 4°C hold. Next, 0.4µL of the mixture of 5µM IgH, Ig? or Igλ chain cloning primers were used for the second round of the PCR in a final volume of 25µL using 0.3µL of DreamTaq DNA Polymerase (5U/µL) (Thermo Fisher), 0.5-2µL of first round PCR product, 0.3µL of 10mM dNTP mix (Thermo Fisher) under the following cycling parameters: 95°C 5 min; 38 cycles of 95°C 30 sec, 55 or 60°C 60 sec, 72°C 1 min, 72°C 10 min and 4°C hold. Then, 5µL of each second round PCR product was loaded in 1.2% agarose gels and run at 100V. Gels were visualized under ultraviolet light, wells with amplified bands were again amplified using second round PCR reaction in 6-tubes of 25µL, run in 1% agarose gels, purified, digested using restriction enzymes and cloned into their respective AbVec vectors (59). Four transformed bacterial colonies were randomly picked, checled for the presence of inserts and sequenced.

### Antibody genes sequence analysis

The sequencing of the antibody genes after their cloning into their corresponding vectors was done commercially from Invitrogen (Thermo Fisher). The sequences were analyzed online through IMGT/V-QUEST (http://www.imgt.org/IMGT_vquest/vquest) and IgBlast (https://www.ncbi.nlm.nih.gov/igblast/). Only selected clones were expressed and purified by the identification of criteria including productive genes, substantial germline mutations (>5%) indicating a history of affinity maturation and long CDRH3 loops.

### Expression of monoclonal antibodies

The HIV-1 monoclonal antibodies AIIMS-P01, 10-1074, and BG18 were expressed in HEK293T (ATCC) or Freestyle 293F cells (Thermo Fisher) by co-transfection of 10µg each of heavy chain and light chain expressing IgG1 plasmids using PEI-Max as transfection reagent. Following 4-6 days of incubation, cells were harvested by centrifugation and filtered through 0.22 mm syringe filter (mdi). The supernatant was added to a Protein A column affinity chromatographic column (Pierce). The column then was washed with 1×PBS and mAbs were eluted with IgG Elution Buffer (Pierce), immediately neutralized with 1M Tris pH 8.0 buffer and extensively dialyzed against 1×PBS at 4°C. After that, mAbs were concentrated using 10kDa Amicon Ultra-15 centrifugal filter units (EMD Millipore), filtered through a 0.22 mm syringe filter (mdi) and stored at −80°C for further use.

### Binding analysis of monoclonal antibodies by ELISA

Briefly, 96-well ELISA plates (Costar) were coated with 5µg/ml recombinant HIV-1 gp120 monomeric proteins overnight at 4°C in 0.1 M NaHCO3 (pH 9.6). Next day, plates were washed thrice with 1×PBS (phosphate buffered saline) and blocked with 15% FBS RPMI and 2% BSA. After 1.5 hours of blocking at 37°C, plates were washed thrice with 1×PBS. Then, serial dilutions of monoclonal antibodies (mAbs) were added and incubated for 1 hour at 37°C. After that, alkaline phosphatase (AP) labelled anti-Fc secondary antibody (Southern Biotech) at 1:2,000 was added and plates were incubated at 37°C for 1 hour. Plates were then washed thrice with 1×PBS and AP substrate tablets (Sigma) dissolved in diethanolamine (DAE) was added and incubated for 30 min at room temperature in the dark and readout was taken at 405nm. The BG505.SOSIP.664 gp140 trimeric ELISA was performed as described previously (6). The polyreactivity ELISA was performed as described previously (60). Briefly, all antigens were coated overnight at 37°C in 96-well ELISA plates (Costar). Next morning, after 3 times washing with 1×PBS the plates were blocked for 1 hour at 37°C 2% BSA and 15% FBS. After again washing the plates 3 times with 1×PBS, the anti-HIV-1 antibodies AIIMS-P01, PGT151 and 4E10 as primary antibodies were added at 5μg/ml diluted in 1×PBS and incubated for 1 hour at 37°C. Next, alkaline phosphatase conjugated anti-human IgG Fc antibody (Southern Biotech) at 1:2,000 dilution in 1×PBS was added after 4 times washing. Readout was taken at 405nm by adding AP substrate (Sigma) in 10% DAE buffer.

### AIIMS-P01 Fab Fragment Preparation

The Fab fragments were generated from 2mg of AIIMS-P01 IgG antibody using a Fab Fragmentation Kit (G Biosciences) according to manufacturer’s protocol. Purity and size of Fab fragments were assessed by SDS-PAGE and binding by surface plasmon resonance (SPR).

### Surface Plasmon Resonance (SPR) analysis

The affinities and binding kinetics of AIIMS-P01 IgG1 and Fab to BG505.SOSIP.664.C2 soluble gp140 trimer and gp120 monomer were determined by SPR on a Biacore T200 (GE Healthcare) at 25°C with buffer HBS-EP+ (10mM HEPES, pH 7.4, 150mM NaCl, 3mM EDTA, and 0.05% surfactant P-20). BG505.SOSIP.664 gp120 trimer and gp120 monomer were first immobilized on a CM5 chip at 600 response units (RU) according to the manufacturer’s standard amine coupling kit protocol (GE Healthcare). The HIV-1 bNAbs AIIMS-P01 and PGT121 were injected at a flow rate of 30µl/min two-fold dilutions starting from 1µM for 300 seconds followed by dissociation for 600 seconds. The cells were regenerated Glycine buffer (pH 2.5). Sensorgrams of the concentration series were corrected with corresponding blank curves and fitted globally with Biacore T200 evaluation software (GE Healthcare) using a 1:1 Langmuir model of binding to calculate the affinity of HIV-1 bNAbs IgG1 and Fab.

### Negative-stain single particle EM and 3D reconstruction

HIV-1 BG505.SOSIP.664 T332N gp140 trimer was incubated with AIIMS P01 IgG and AIIMS P01 Fab at a different concentration ratio of 1:1, 1:2 and 1:3 for one hour at 4°C. The samples were prepared using the conventional negative staining methods (61). Briefly, 3.5 µl of samples were adsorbed on UV treated carbon-coated copper grids for one minute. This was followed by washing with three drops of water and staining with 2% uranyl acetate for 20 seconds. Samples were imaged at room temperature by using a Tecnai T12 electron microscope equipped with the LaB6 filament at 120 kV accelerating voltage. The images were recorded at a magnification of 75k on a side-mounted Olympus VELITA (2Kx2K) CCD camera. 100 tilt pair images were collected at 0° and 45° tilt angle by tilting the sample stage and all the data were collected at a defocus range of ∽-1.5 to −1.8µm with the final pixel size of 2.0 Å at the specimen level. Particles were excised from the untilted images automatically using “swarm” option of e2boxer.py in EMAN 2.12 suite (62). Total 11243 particles were selected from untilted micrographs. The reference free 2D class averaging was calculated using simple_prime2D in SIMPLE 2.1 software package (63) and the dataset was classified into 120 classes. For an initial model generation, we employed the random conical tilt pair (64) image reconstruction techniques to determine the back-projected 3D model of HIV-1-Fab complex (HIV-1 BG505.SOSIP.664-T332N gp140 with AIIMS-P-01-Fab) from the tilted dataset using XMIPP (X-windows based microscopy image processing package (65). Simultaneously, an initial model was calculated from reference free class averages using e2initialmodel.py in EMAN 2.12 suite. Both the initial models are structurally similar, and used as an initial model for further data processing. The initial map was filtered to 40 Å using RELION-1.4 (66) and used as a reference model for 3D classification. About 7640 particles were selected from 3D classification where Fab is clearly visible. These particle sets were used for final auto refinement in Relion 1.4 software package. The resolution for the map was calculated using the 0.5 cutoff of Fourier shell correlation and estimated resolution is 26Å. Three-dimensional EM map was visualized by UCSF Chimera (67). The crystal structure of Fab (3PIQ) and HIV trimer (4NCO) were docked into the EM map using the automatic fitting option of Chimera software package.

### Competitive ELISA

The competitive ELISAs were performed as described previously (12). The ELISA plates were coated overnight at 4°C with 10μg/ml of anti-C5 (D-7324) antibody (Aalto Biosciences) in 0.1 M NaHCO3 (pH 9.6). Next day, plates were washed thrice with 1×PBS blocked for 1 hour at 37°C with 2% BSA and 15% FBS. Then, 1:4 serial dilution of competing antibodies were added with a starting concentration of 40µg/ml in the presence of biotinylated AIIMS-P01 at a fixed concentration of 5µg/ml. Then, plates were washed thrice and streptavidin-HRP was added at (1:1000) dilution in 1×PBS followed by addition of 3,3′,5,5′-Tetramethylbenzidine (TMB) substrate (Biolegend). The reaction was stopped by adding H_2_SO_4_ stop solution before measuring binding at 405nm. These experiments were performed twice.

### Cell surface binding assay

The binding of the antibodies to cell-surface HIV-1 envelopes was assessed using a flow cytometry-based assay as described earlier (60). HEK293T cells (2×10^5^ cells) were transfected with 5 µg of HIV-1 envelope expression plasmid DNA using PEI-Max transfection reagent followed by harvesting at 48 hours post-transfection and incubated at 5µg/ml concentration of antibodies for 1 hour at 37°C. After washing with 1×PBS, cells were next stained with phycoerythrin (PE) conjugated anti-human IgG Fc antibody (Biolegend) for half an hour at room temperature followed by acquiring the data by flow cytometry using a BD FACS Canto II. Data were analyzed using FACSDiva software. Percent binding was calculated as the percentage of PE-positive cells with the background (antibodies binding to mock-transfected cells) subtracted. Analyses were performed in GraphPad Prism 5.0.

### HIV-1 envelope amplification and cloning

The HIV-1 enveloped AIIMS_330 autologous pseudoviruses were generated by single genome amplification (SGA) as described previously (45). Briefly, viral RNA was extracted from 140µl of AIIMS_330 plasma of each independent time point by using the QIAamp Viral RNA Mini Kit (Qiagen) followed by cDNA synthesis using Superscript III reverse transcriptase cDNA synthesis kit (Invitrogen) as per manufacturer provided protocol. The HIV-1 envelope (rev/env cassette) was amplified by nested PCR using primers and reaction conditions as described earlier (45). The amplicons were sequenced commercially from Eurofins Genomics. The SGA amplicons with full-length envelope genes were molecularly cloned into the pcDNA3.1 Directional Topo vector (Invitrogen), transformed into TOP10 competent cells and sequenced.

### ADCC assay

The ADCC assays were done as previously described (68) with few modifications. The 5×10^7^ target cells (CEM.NKR-CCR5-sLTR-Luc cells) (NIH AIDS reagent program) were infected with HIV-1 NL4-3, HIV-1 JRFL, or MuLV at 200 TCID50 by incubating at 37°C for 1 hour in a final volume of 200µl. Two days post infection, the infected target cells were washed three times with RPMI medium and incubated in round bottom 96-well plates with effector cells (KHYG-1 cell line) expressing CD16 at a 10:1 E:T ratio in the presence of serial dilutions of AIIMS_330 and healthy donor plasma, AIIMS-P01 and VRC01 HIV-1 bNAb. After an 8 hr incubation, cells were lysed with luciferase substrate (BriteGlo, Promega), and luciferase activity in relative light units (RLUs) was measured with a Tecan multi-plate mode reader. The ADCC activity of the plasma and antibodies were calculated from the mean RLU for triplicate wells at each antibody concentration/plasma dilution relative to the means for background and maximal RLUs from replicate wells containing uninfected and infected cells, respectively, incubated with NK cells but without antibody.

### Quantification and statistical analysis

All statistical analysis was done with GraphPad Prism software version 5.

### Data availability

The sequences of the AIIMS-P01 heavy chain and light chain variable region have been deposited in GenBank with accession numbers MH267797 and MH267798 respectively. The sequences of the AIIMS_330 autologous viruses have been deposited in GenBank and their accession numbers are pending. The accession number for the EM map of AIIMS-P01 Fab in complexed with BG505.SOSIP.664.C2 T332N gp140 trimer reported in this study is EMD: 6967.

## Acknowledgments

This antibody work was supported by Science and Engineering Research Board (SERB), Department of Science and Technology (DST), India grant (EMR/2015/001276). The HIV-1 pseudoviruses generation work was supported by (BT/PR5066/MED/1582/2012) Department of Biotechnology (DBT), India. The EM work and consumables were supported by (SERB-EMR/2016/005407 & BT/INF/22/SP22844/2017). The senior Research Fellowship to SK was supported by SERB, India. We are very much thankful to pediatric donor AIIMS_330 for participating. We are very much thankful to NIH AIDS reagent program for HIV-1 research reagents, Neutralizing antibody consortium (NAC), IAVI, USA for HIV-1 neutralizing antibodies; Prof. Michel C. Nussenzweig, Rockefeller University, USA for 10-1074 and BG18 antibody expression plasmids; Prof. Dennis R. Burton, The Scripps Research Institute, USA, Prof. John P. Moore & Prof. Rogier Sanders, Cornell University, USA and University of Amsterdam, Netherlands, respectively for BG505.SOSIP.T332N.664 plasmids; Prof. Lynn Morris, NICD, SA for N160 and N332 mutant plasmids; Prof. Patrick C. Wilson, University of Chicago, USA for AbVec antibody expression plasmids; Dr. Jacob Kopicinski, Imperial College London, UK for helping in designing the FACS antibody panel; Dr. Dinakar Salunke, Dr. Pawan Malhotra, Naresh Sahoo, ICGEB, New Delhi, India, for ICGEB SPR Facility, Dr. Likhesh Sharma (GE Healthcare) for helping with the SPR experiments and Pradeep Kumar, BD Biosciences, India, for his assistance in sorting of single B cells by FACS.

## Declaration of Interests

K.L., S.K., and R.L. have a pending Indian provisional patent application for pediatric bNAb AIIMS-P01 described in the present study. All the authors have read and approved the manuscript for publication and declare no competing interests.

## Author Contributions

S.K. planned and performed the experiments, analyzed the data, and wrote the original manuscript; H.P. helped S.K. in standardizing amplification protocols for antibody genes; M.A.M. and N.M. generated AIIMS_330 autologous pseudoviruses, N.M. analyzed the FACS data and helped S.K. in HIV-1 pseudoviruses generation and neutralization assays; H.A. helped S.K. in Flow cytometry experiments; R.L. and S.K.K. provided the donor sample, were responsible for clinical care of the patient; E.S.R. helped S.K. in size-exclusion chromatography experiments; A.C. provided the scientific inputs for single B cell sorting technique and reviewed the manuscript; S.D. performed and solved the 3D structure from negative stain EM experiments, along with H.A.S. and reviewed the manuscript; and K.L. conceived, planned and supervised the experiments, reviewed, edited and finalized the manuscript. All authors reviewed the manuscript and approved the final version of the manuscript.

